# Spatio-temporal model and machine learning method reveal process of phenological shift under climate change of North Pacific spiny dogfish

**DOI:** 10.1101/2022.03.01.482455

**Authors:** Yuki Kanamori, Toshikazu Yano, Hiroshi Okamura, Yuta Yagi

## Abstract

Climate change has disrupted natural phenological patterns, including migration. Despite extensive studies of phenological shifts in migration by climate change and driving factors of migration, a few issues remain unresolved. In particular, little is known about the complex effects of driving factors on migration with interactions and nonlinearity, and partitioning of the effects of factors into spatial, temporal, and spatio-temporal effects. The Pacific spiny dogfish *Squalus suckleyi* (hereafter “spiny dogfish”) is a coastal elasmobranchii that migrates southward for parturition and northward for feeding in the western North Pacific. Here, to elucidate the migration patterns as well as the driving factors under climate change, we first examined long-term changes in the timing and geographic location of migration by applying a spatio-temporal model to ca. 5-decade time series data for the presence/absence of spiny dogfish in the western North Pacific. We then evaluated the spatial, temporal, and spatio-temporal effects of driving factors (sea surface temperature [SST], depth, productivity, and magnetic fields) on seasonal occurrence patterns using a machine learning model. We found that the migration area did not change over ca. 5 decades, whereas the migration timing advanced by a month after 2000. The spatial effects of magnetic fields and depth were consistently large and the spatial and spatio-temporal effects of SST increased in the migration season, even though temporal effect of SST was always weak. These results suggest that the migration area of spiny dogfish was stable over time because their spatial distribution was determined by geographic features, whereas the migration timing advanced by tracking a suitable SST location which increased steeply after 2000. Therefore, temperature as well as other factors influence migration simultaneously under climate change and underline the importance of paying attention biotic/abiotic factors including temperature and process-based understanding to predict future impacts of climate change on phenological shifts.

## 1 Introduction

Climate change has substantially affected organisms worldwide (e.g., Walther et al. 2002, Hoegh-Guldberg and Bruno 2010, Walther 2010, Pecl et al. 2017). Changes in phenology (i.e., seasonal timing of life-history events, such as migration and reproduction) are thought to be an indicator of the effect of climate change on organisms since temperature, which has changed under climate change, is a predominant factor affecting phenology. Recent meta-analyses have revealed the magnitude of phenological shifts relative to a historical baseline (e.g., Parmesan and Yohe 2003, Root et al. 2003, Menzel et al. 2006, Parmesan 2007, Poloczanska et al. 2013, Asch 2015) and variability in the magnitude and patterns among taxa, ecological traits, and ecosystems (e.g., Edward and Richardson 2004, Burrows et al. 2011, Poloczanska et al. 2016, Brown et al. 2016). However, studies about phenological shifts often have gaps in terms climate change and ecology. Since species are affected by various, complex external factors with complex interactions and nonlinear effects (e.g., Araújo and Guisan 2006, Guisan et al. 2006, Sala et al. 2000, Sugihara et al. 2012, Ye et al. 2015, Ryo et al. 2018, Nakayama et al. 2020), phenological shifts should be related to complex interactions and nonlinear effects; simple statistical methods such as correlation, linear regression, and similarity may lead to erroneous results.

Migration, which is a phenological event, plays a role in population and ecosystem dynamics as well as in ecosystem services such as tourism (e.g., Gallagher and Hammerschlag 2011, Bauer and Hoye 2014, Teitelbaum and Mueller 2019).

Recent developed tracking technology has provided insight into the drivers of migration. One major driver of migration is ecological factors such as prey (e.g., Mathot et al. 2007, Sherrill-Mix et al. 2008, Milner-Gulland et al. 2011, La Sorte and Graham 2021) and temperature (e.g., Marra et al. 2005, Sherrill-Mix et al. 2008, Kanamori et al. 2019, Archer et al. 2020). This type of factors leads to migration through changes in physiological conditions such as reproduction; migration itself also affects physiological conditions, thereby regulating the timing and location of migration. Another major driver of migration is wayfinding factors such as magnetic fields (e.g., Wiltschko and Wiltschko 1995, 2005, Diebel et al. 2000, Cain et al. 2005, Aderson et al. 2017, Keller et al. 2021) and olfactory cues (e.g., Koch et al. 1969, Bett and Hinch 2015, Bett et al. 2018, Durif et al. 2021), which regulate the route and accuracy of migration. Despite extensive evidence and predictions that these factors relate to migration, some challenging problems remain unresolved. The first problem involves considering the complex effects of driving factors on migration because species generally has been affected by various factors with interactions (e.g., Araújo and Guisan 2006, Guisan et al. 2006, Sala et al. 2000, Ryo et al. 2018) and nonlinearity effects (e.g., Sugihara et al. 2012, Ye et al. 2015, Nakayama et al. 2020, Satterthwaite et al. 2020). The complex descriptions of the relationship between species and migration-driving factors make it difficult to interpret the results of analyses; therefore, a method for easily interpreting the results is needed. The second problem involves partitioning the effects of factors into spatial, temporal, and spatio-temporal effects. For example, we consider a general case in which we use time-series data of a species within a given spatial range and use a generalized linear model to analyze the relationship between species occurrence and a factor such as temperature. Because the data include both temporal and spatial information, the interpretation of the effect of the factor on species occurrence (i.e., regression coefficient) is vague. In other words, does the factor affect the variation in species occurrence spatially, temporally, or both? Use of this approach can lead to further understanding of migration, including whether species actively seek out specific conditions or merely react to changes in conditions (Schlaff et al. 2014).

Elasmobranchs (sharks, skates, and rays) are apex predators with long-distance migration in marine systems (Heupel et al. 2015). Their migration is thought to play an important role in marine communities and ecosystems by horizontally and temporally connecting habitats and transferring energy via the trophic cascade (e.g., Dulvy et al. 2000, Heithaus et al. 2008, Polovina et al. 2009, Heithaus et al. 2012, Hummerschlag et al. 2019). Although this migration also underpins their populations through gaining energy, they have been threatened with extinction (Worm et al. 2013, Dulvy et al. 2014) due to overfishing (e.g., Dulvy et al. 2021, Pacoureau et al. 2021) and climate change (e.g., Chin et al. 2010, Niella et al. 2021, Hammerschlag et al. 2022). Hence, it is necessary to elucidate the spatio-temporal changes in their migration patterns and driving factors for the development of effective conservation and management strategies. Many studies have shown that ecological factors, such as temperature and prey as well as other drivers (e.g., salinity, pH, and tide), trigger the migration (e.g., Ackerman et al. 2000, Torres et al. 2006, Carlisle and Starr 2009, Ortega et al. 2009, Ubeda et al. 2009, Niella et al. 2021). In addition, elasmobranchs are known as species who use magnetic field when migration. Indeed, they perceive magnetic fields (e.g., Kalmijin 1982, Meyer et al. 2005, Mora et al. 2014, Anderson et al. 2017, Newton and Kajiura 2020) and a shark showed that the ability to perceive magnetic fields is used for navigation (Keller et al. 2021).

The North Pacific spiny dogfish *Squalus suckleyi* (hereafter “spiny dogfish”) is a demersal and coastal shark, which had less attention in the previous studies of migration of sharks. They widely distribute in the North Pacific, from the Bering Sea to the coast of Japan and from the Gulf of Alaska to California (Orlov et al. 2012). In the western North Pacific, they migrate southward from October to December for parturition, after which the adults and their offspring migrate northward around April for feeding (Kojima 1958; Fig. 1). It is possible that they migrate by following changes in water temperature (Kojima 1958, Ebert 2003). Their preferred water temperature is 6°C to 10°C in the western North Pacific (Yano et al. 2017). A previous study suggested that they may be opportunistic feeders with no specific target prey (Mecklenburg et al. 2018), while another study indicated that their distribution may be related to the presence of potential prey, including sardine, walleye pollock, and squid (Yano et al. 2017). They are a commercially important target species in Japan, the United States, and Canada (Orlov et al. 2012). In Japan, they have been caught in the Sea of Japan and in the western North Pacific. Their stock assessment has been conducted by two abundance indices (catch per unit effort) from the bottom trawl fisheries data in the western North Pacific and the bottom longline data in the Tsugaru Strait (Fig. 1).

**Fig. 1.**
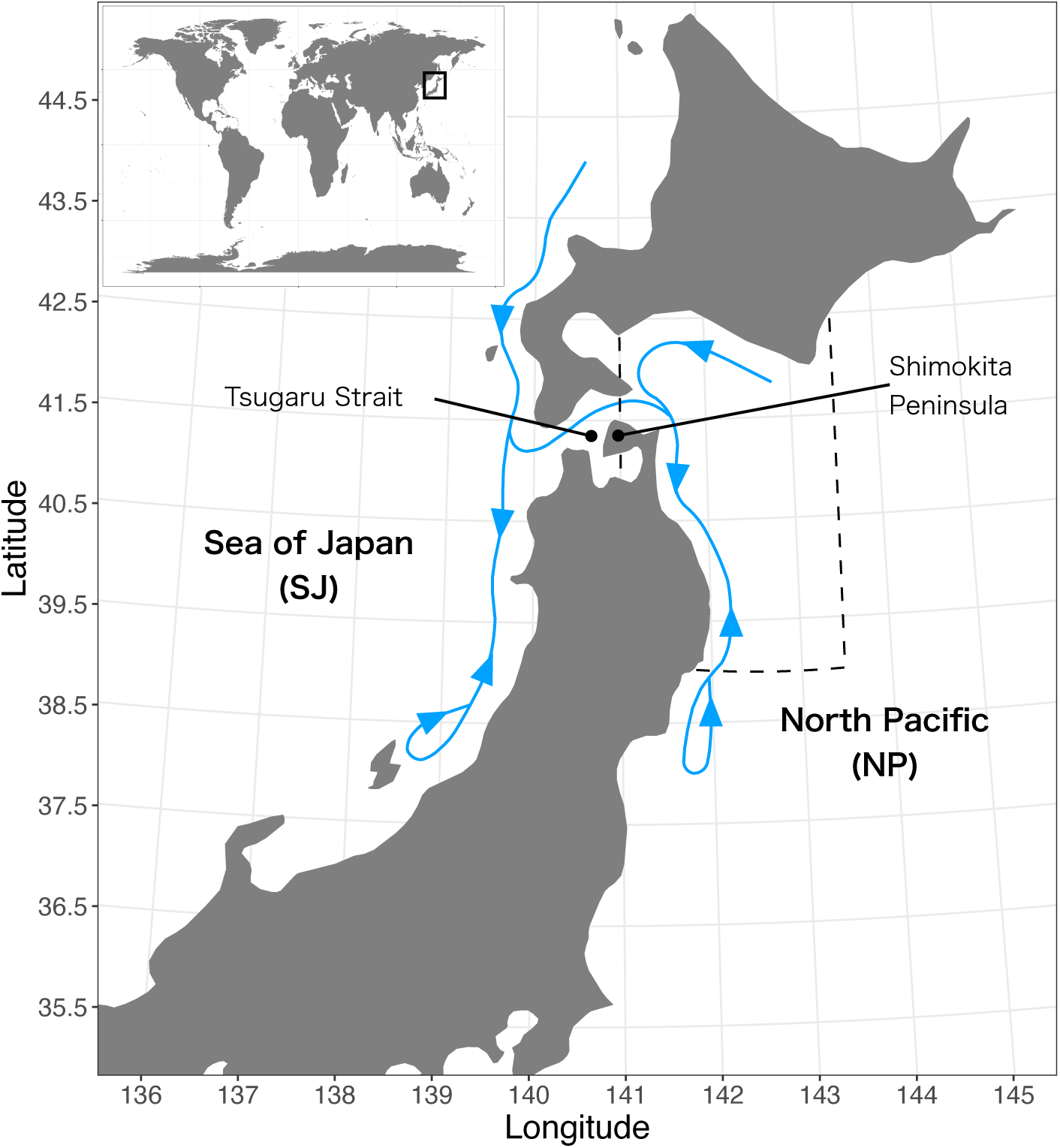
Study area. North Pacific spiny dogfish *Squalus suckleyi* starts migrating southward from October to December for parturition and then migrates northward around April. In this study, the Sea of Japan (SJ) and the North Pacific (NP) were defined as the areas west and east of 140.5°E, respectively. Stock assessment in Japan has been conducted using bottom longline data from the Tsugaru Strait and bottom trawl data within the dashed area

Here, to elucidate the phenological shift in migration and its processes under climate change, we first examined the long-term changes in timing and geographic location of migration by applying a spatio–temporal model to ca. 5-decade time series data for the presence/absence of spiny dogfish in the western North Pacific. To understand the processes of the change in migration, we then evaluated the spatial, temporal, and spatio-temporal effects of driving factors (productivity, sea surface temperature [SST], and depth, and magnetic fields) on seasonal occurrence of spiny dogfish using a machine learning model and an interpretable machine learning technique.

## 2 Materials and Methods

### 2.1 Data sets

#### 2.1.1 Occurrence data for spiny dogfish

Daily catch statistics for bottom trawl fisheries from 1972 to 2019, except for the fishery closures season (July and August), was used. The spatial resolution was 10′ latitude *×* 10′ longitude horizontal square in the area from 138°E to 142°E and 37°N to 42°N. We defined the area west of 140.5°E as SJ (Sea of Japan) and the area east of 140.5°E as NP (North Pacific) (Fig. 1).

The daily catch statistics included date, geographic location (longitude and latitude), fishing effort (number of tows), catch weight in each fish category including spiny dogfish, and so on. Even though the data had geographic location, the location was not all fishing points in a day but only one location in a day; fishermen recorded the total catch weight in each fish category and geographic location of the most caught fish category every day. Accordingly, there is the possibility that record of catch weight of spiny dogfish was low or high in a point where spiny dogfish inhabits or doesn’t inhabit. Therefore, to estimate the monthly distribution in each year, we transformed the catch weight of spiny dogfish into a binary variable (presence/absence) taking a value of 1 when catch weight > 0 and 0 otherwise.

Spiny dogfish was assigned to the category elasmobranchs in SJ from 1972 to 2019 and in NP from 1972 to 1993; therefore, the catch weight may include the catch weight of other elasmobranchs. However, we have not simultaneously caught both spiny dogfish and other elasmobranchs at the same sites during bottom trawl surveys in SJ for stock assessments (personal observation). Moreover, Yano et al. (2022) showed evidence that data for elasmobranchs in SJ and NP mostly reflect the catch weight of spiny dogfish.

#### 2.1.2 Data collection for candidate driving factors

Data for three ecological factors (productivity, SST, and depth) and one wayfinding factor (magnetic fields) were collected.

##### Productivity

Productivity was defined as the total catch weight of all fish categories other than spiny dogfish in a month divided by the total efforts in a month using the daily catch statistics. We assumed that as the total catch weight increases, the abundance of prey that spiny dogfish can use increases. The productivity data were aggregated over two spatial resolutions (0.5° latitude and 1.5° latitude) owing to uncertainty regarding how spatial scale affects the occurrence of spiny dogfish.

##### SST

The mean global daily SST since 1982 with 25′ latitude × 25′ longitude horizontal square resolution in the study area archived as MGDSST by the Japan Meteorological Agency (https://ds.data.jma.go.jp/gmd/goos/data/database.html) was collected. The monthly mean SST exhibited positive trends, and the trends increased after 2000 from September to June (Fig. S1).

We calculated both the monthly mean SST and monthly coefficient of variation (CV) in each year and each data point of MGDSST data since climate change generally increases both the mean and variability of temperatures. Then, both the mean and CV of SST were given for the nearest predicted points with spiny dogfish occurrence using a spatio-temporal model (see 2.2.1 for details) because the points of MGDSST data and the predicted points were distinct. The mean, minimum, and maximum distances between the points of MGDSST data points and the predicted points were ca. 5.8 km, 0.9 km, and 11 km, respectively. The mean and CV of SST before 1981 were not available because MGDSST was recorded from 1982.

##### Depth

The depth data in the study area were collected from the Japan Oceanographic Data Center (https://ds.data.jma.go.jp/gmd/goos/data/database.html).

In the study area, the depth averagely changes with longitude. For the area within ±5′ latitude from a predicted point, depth data for less than 1 km were obtained. After that, mean depth and coefficient of variation (CV) which means drastic changes in depth around predicted points (i.e., submarine topography) were calculated. Only three predicted points did not have the depth data within 1 km and ±5′ latitude; for these three predicted points, depth data within 3 km were obtained.

##### Magnetic fields

Yearly values of three parameters (declination, inclination, and total intensity; these three parameters are a combination showing magnetic fields) were calculated in each predicted point using the gufm1 and IGRF models developed by the National Oceanic and Atmospheric Administration (https://www.ngdc.noaa.gov/geomag/).

### 2.2 Data analyses

#### 2.2.1 Estimation of spatio-temporal variation in occurrence

To estimate spatio-temporal variation in the monthly occurrence of spiny dogfish, we used the barrier model (Bakka et al. 2019) which is a state-of-the-art spatio-temporal model. This model accounts for spatio-temproal changes in fishing location, which likely cause estimation biases (Thorson et al. 2017a). Additionally, this model accounts for spatial autocorrelation in the physical barrier, which is necessary because the study area contained some islands with complex coastline (Bi et al. 2020). The occurrence rate *p*_*i*_ for each sample *i* was approximated using a logit-linked linear predictor as follows:

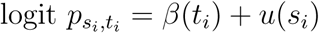

where *β*(*t*_*i*_) is the effect of time *t* defined as a year-month combination (i.e., January 1972 is *t* = 1, February 1972 is *t* = 2, …, December 2020 is *t* = 480), which was estimated as a first-order random walk due to the lack of data for the fishery closures season (July and August). *u*(*s*_*i*_) is the spatial random effect for location *s* in Gaussian random field with a Matérn covariance function through stochastic partial differential equations (SPDE). Although the full conditional distribution at the site in two-dimensional space has the expectation using small set of neighbors’ under the Matérn model through SPDE, the paths that crossed the physical barrier (e.g., land and coastal line) are eliminated in the barrier model by adding the idea of Simultaneous Autoregressive model into Matérn model through SPDE. Thus, *u*(*s*), which is a spatial random effect in the barrier model, is a solution to the following SPDE using the finite element method:

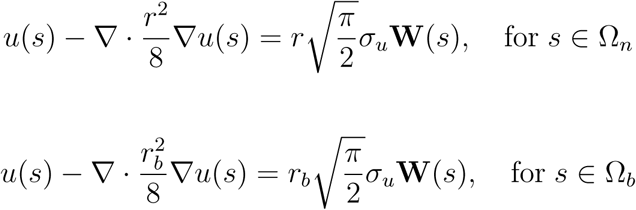

where *u*(*s*), *s* ∈ Ω ⊆ ℝ^2^ is the Gaussian random field, *r* and *σ*_*u*_ are constants, 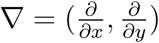, and ***W*** (*s*) denotes spatial white noise. Ω_*n*_ is the normal area, Ω_*b*_ is land, and their disjoint union gives the whole study area Ω. For further details, see Bakka et al. (2019).

As described above, geographic locations included uncertainties because only one geographic location was not recorded in a day for each fishermen (see details for *2*.*1*.*1 Occurrence data for spiny dogfish*). If covariates were included in this model, occurrence rates would be estimated based on inaccurate relationships between the occurrence pattern and covariates. Thus, we did not include any covariates in this model.

Parameters in the model were estimated by integrated nested Laplace approximation algorithm (Rue et al. 2009) using the R-INLA package (Bakka et al. 2018) in R 3.6.3 (R Development Core Team 2021). Then, we predicted the occurrence rates 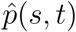 in the study areas. Hereafter, we call the points with predicted occurrence rates as “predicted points.” The distances between the predicted points were ca. 10 km.

#### 2.2.2 Migration patterns

##### Geographic location

To evaluate long-term changes in the geographic location of migration, we used two measurements. First, the mean spatial distribution of spiny dogfish in each month was used to find the average migration patterns over 5 decades because it was difficult to find the patterns from the extensive number of maps of predicted occurrence rates. The unimodality in each predicted point and month was checked by Silverman ‘s test which was conducted using the package *silvermantest* in R. The raw maps using the predicted occurrence rates 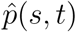 are shown in the Appendix (Figs. S2-7).

Second, we used the inter-annual patterns of the center of gravity (COG) in each month using the predicted occurrence rate for each predicted point 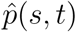. The latitudinal and longitudinal COG at time *t*, COG_Lat._(*t*) and COG_Lon._(*t*) were estimated as:

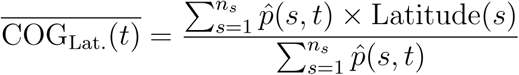

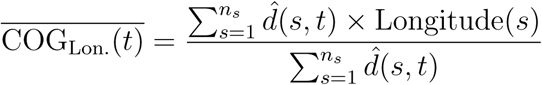

where Latitude(*s*) and Longitude(*s*) are the latitude and longitude of location *s*. We estimated COGs in both SJ and NP because the study area was separated by land and a single COG was not appropriate. The changes in COGs were small in both of SJ and NP, thus the line graphs using the values of COGs are shown (Fig. S8).

##### Timing

To examine the long-term changes in the timing of migration of spiny dogfish, the peak of migration in each year was estimated in both JS and NP. We first calculated the monthly mean occurrence rates in both SJ and NP using estimated occurrence rates 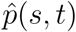 to obtain the migration patterns in each year (Figs. S9). We then defined the mode of the migration patterns as the peak of migration. Here, the “year” was defined as September of year *t* to June of *t* + 1 because spiny dogfish begins to migrate southward in autumn and data were not available for July and August due to fishery closures season.

In some cases, unimodality was not supported by Silverman ‘s test, especially for migration patterns in NP. For example, a histogram of 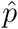 in September 1972 in NP was not unimodal, with high estimated occurrence rates around the Shimokita, indicating that the mean value is not representative for analyses of migration patterns in each year. Therefore, we also obtained the migration patterns in each year using the mode and median for sensitivity analyses (Figs. S10-11). Then, we defined the mode of the migration patterns in each year as the timing of migration (Figs. S12-13), similar to the analysis using mean values. The results for long-term changes in the timing of migration were consistent, although there were slight differences between results obtained using different metrics, particularly between the mean and the mode (Figs. S12-13).

#### 2.2.3 Evaluating the spatial, temporal, and spatio-temporal effects of driving factors

To model the relationship between the estimated occurrence rate of spiny dogfish and driving factors, we used the gradient boosting method which is based on machine learning algorithm (e.g., De’ath 2009, Hastie et al. 2009, Maloney et al. 2012, McLaren et al. 2018, Shiferaw et al. 2019). The method sequentially generates large numbers of weak learners and finally combines them. This combination of weak learners can describe nonlinearity between a response variable and covariates as well as complex interactions among covariates.

Firstly, we checked correlations among covariances in each month. The variables related to the magnetic fields (declination, inclination, and total intensity) were consistently correlated with each other. Although this presents an issue when interpreting the model, we did not eliminate any magnetic fields variables; instead, we resolved the issue by the interpretation method described in the last paragraph in this section. Strong correlations among other driving factors were not detected. Then, after data for each month were divided into training data (70%) and test data (30%), we tuned seven hyperparameters (nrounds, max depth, eta, gamma, colsample bytree, min child weight, and subsample) and trained the models by ten-fold cross-validation using the training data. This tuning was conducted using the packages *caret* (Kuhn 2008, Kuhn 2020) in R. The hyperparameters were obtained by five grid searches and the optimal models (i.e., high performance models) were chosen based on the root mean squared error (RMSE). All of the driving factors were scaled and standardized. This analysis was conducted with both high-resolution and low-resolution productivity data. We determined the best model based on the RMSE when using the test data. The model using the productivity with low-resolution was selected as the best model in all months; accordingly, the results using productivity with low-resolution are presented.

To demonstrate that the gradient boosting model had a higher predictive value than that of a linear model, we also analyzed the relationship between the estimated occurrence rate and driving factors for each month using the training data by a linear model which included the terms of all driving factors, pairwise second-order interactions, squared mean SST and mean depth. We then compared between RMSE of gradient boosting and the linear model when using the test data.

To evaluate the spatial, temporal, and spatio-temporal effects of driving factors on migration, we used grouped permutation feature importance (GPFI). GPFI measures the change in prediction errors between the best model and when permuting a “single set of covariates.” A greater increase in the prediction error is interpreted as more important covariate for the prediction (Molnar 2020). The original idea of this method is to calculate the importances of each covariate by permuting correlated covariates as a single set since like a linear model, the calculated importance when permuting a single covariate which correlated other covariances is unstable and difficult to interpret. We applied this method and calculated the RMSE as the prediction error when permuting within spatial, temporal, spatio-temporal for each covariate. We then calculated the *L*_2_ norm between the RMSE from the best model and the RMSE from the permutations to evaluate the spatial, temporal, and spatio-temporal effects of driving factors. For example, we consider data for 10 sites and 10 years for a species and a factor (e.g., temperature). The *L*_2_ norm between the RMSE from permuting a factor in a fixed site (e.g., site 1) and the RMSE from the best model indicates the importance of the temporal effect of the covariate because the factor data are permuted among years. This permutation was conducted 100 times in each month, and covariates related to magnetic fields (declination, inclination, and total intensity) were treated as a single set due to the observed correlations. We can grasp the macroscopic patterns from figures that summarize the results for all months (Figs. S14-18).

## 3 Results

### 3.1 Long-term changes in migration patterns

The spatial distribution of spiny dogfish clearly changed seasonally (Figs. 2 and S2-7). Although the mean occurrence rates were low in the whole study area from September to December, spiny dogfish occurred on the north side of the Shimokita Peninsula. In January and February, the mean occurrence rates increased significantly around the Shimokita Peninsula and increased on the north side of SJ and along the coastline of NP. From March to May, the mean occurrence rates in SJ and NP decreased gradually, although the occurrence rates on the north side of the Shimokita Peninsula remained slightly elevated. In June, the mean occurrence rates were somewhat high only on the north side of the Shimokita Peninsula.

**Fig. 2.**
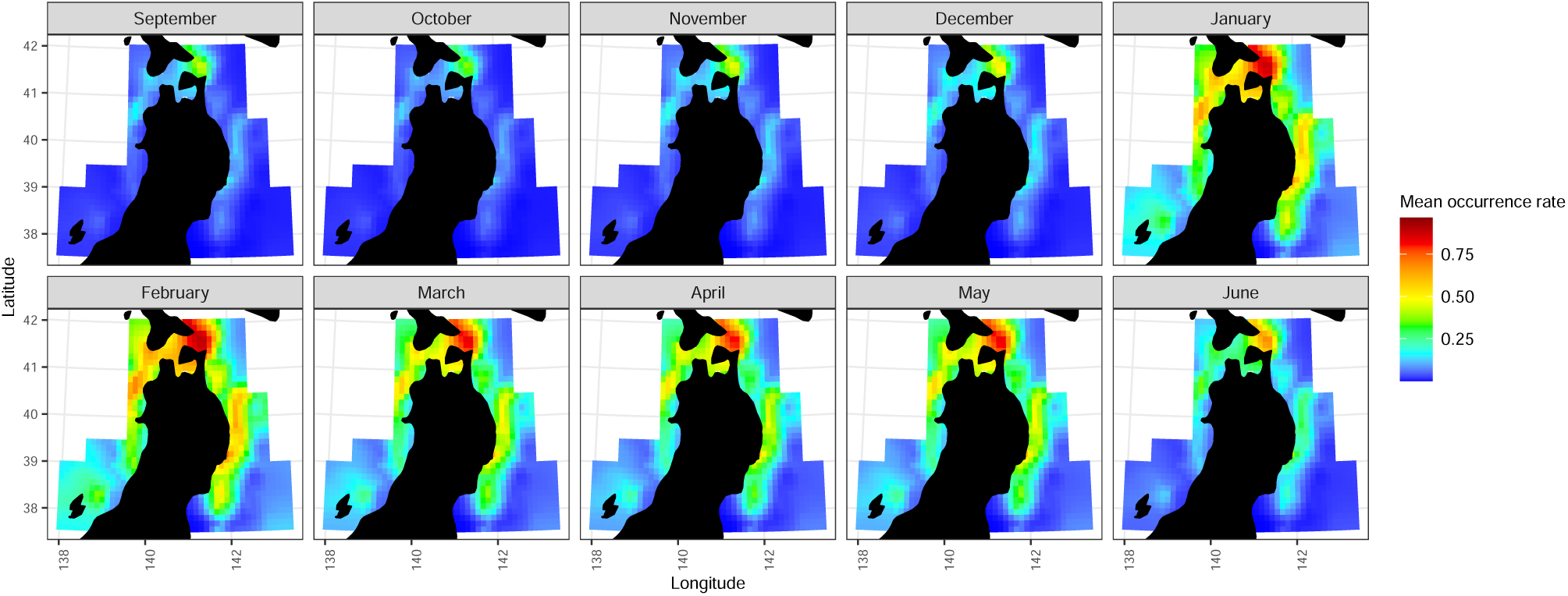
Mean spatial distribution of spiny dogfish from September to June, as estimated using the model

COG did not change in any month during the ca. 5-decade period (Fig. 3). The longitudes of COG in SJ and PO did not change in any month. There were greater fluctuations in latitudes of COG in SJ and NP than in longitudes (Fig. S8). Although the latitude tended to decrease from 1972 to around 2015, the magnitude of change was quite small.

**Fig. 3.**
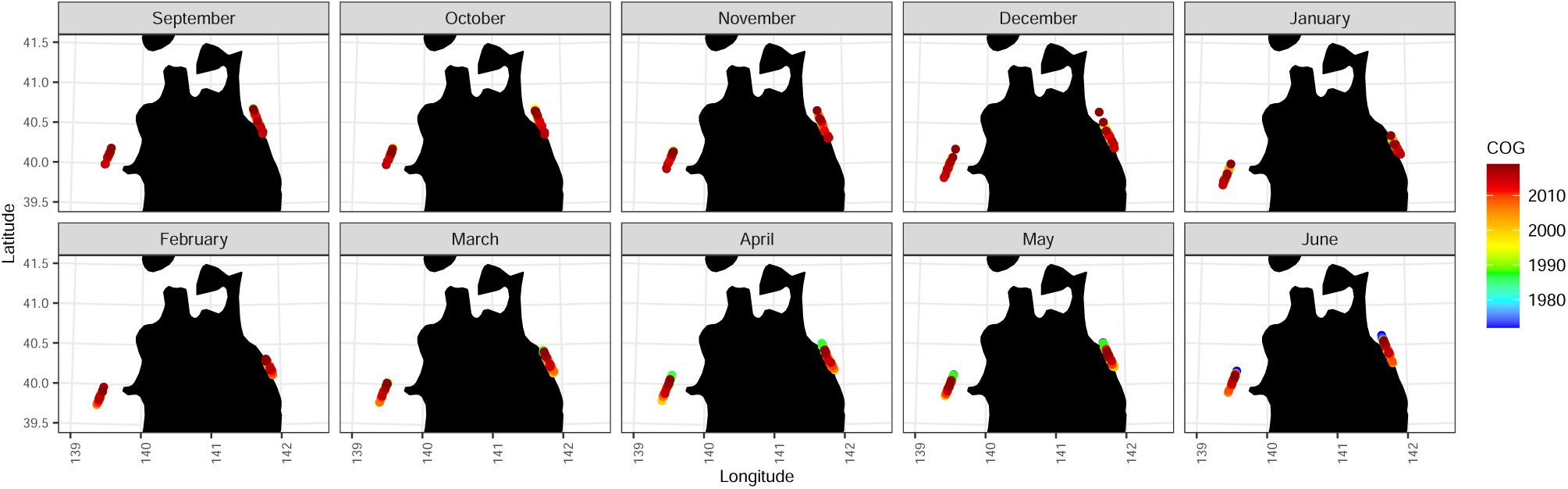
Long-term changes in the center of gravity (COG) from September to June

The timing of migration advanced in both SJ and NP (Fig. 4). Before 2000, migration was observed mainly in February, compared with January after 2000. Consistent results were obtained when using the mode and median of estimated occurrence rates (Figs. S12-13).

**Fig. 4.**
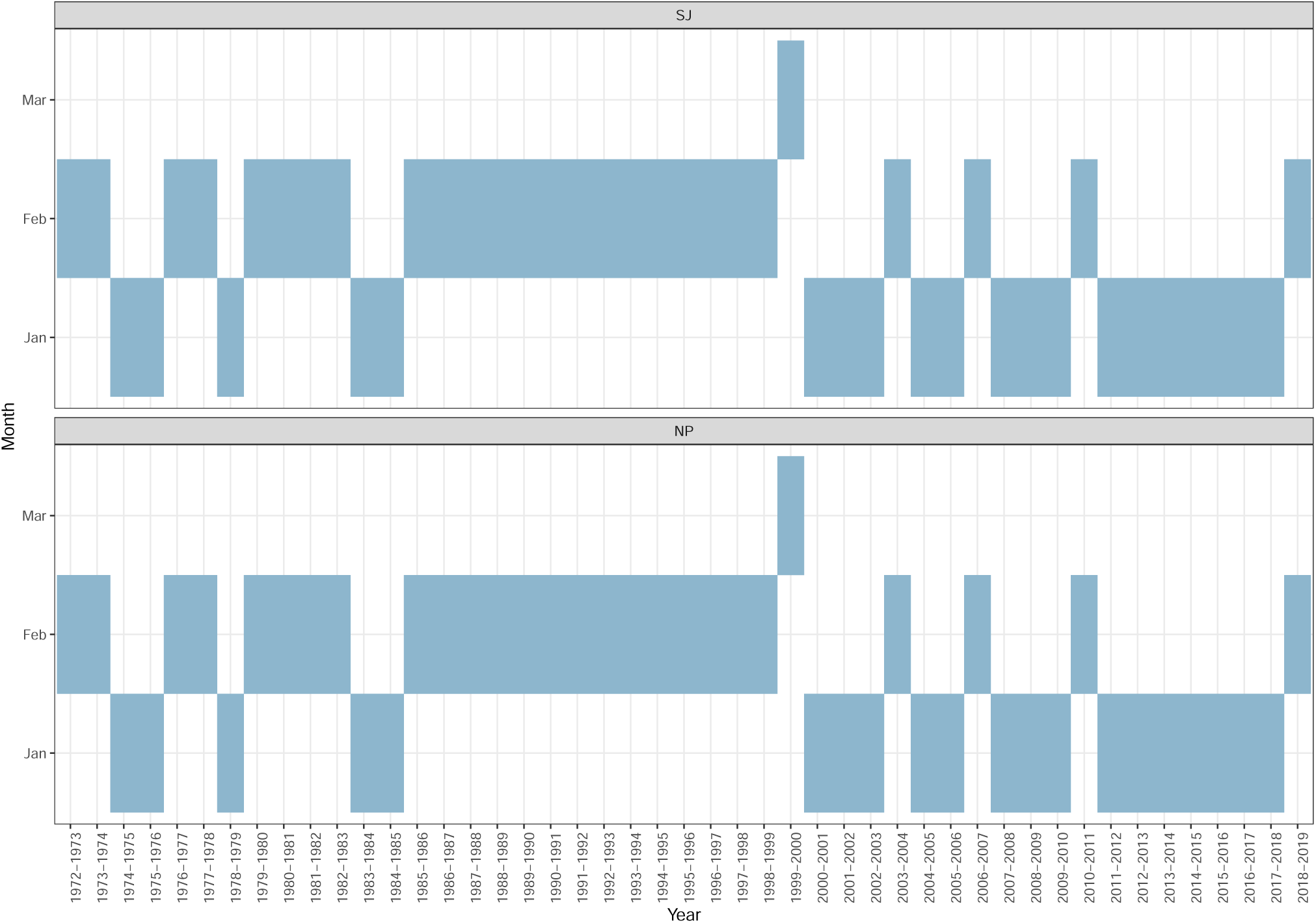
Long-term changes in the timing of migration of spiny dogfish in the Sea of Japan (SJ) and in the western North Pacific (NP). Here, each year started from September to June because spiny dogfish start to migrate in autumn

### 3.2 Spatial, temporal, and spatio-temporal effects of driving factors on migration

All gradient boosting models had always higher predictability comparing with the linear models, although the linear models were slightly complex due to including interactions among variables and square terms for nonlinearity (Tables S1-2).

The effects of driving factors, particularly SST and productivity, changed seasonally (Fig. 5). The spatial effects of magnetic fields and depth and the spatio-temporal effect of magnetic fields were always large (shown in a pink boxplots). Following those factors, the spatial, temporal, and spatio-temporal effects of productivity, the spatial effects of CV of depth, and the spatial and spatio-temporal effects of SST also tended to be large (the yellow box plots). The spatial and spatio-temporal effects of SST tended to increase in February and then decrease in April, whereas the effects of productivity tended to decrease in February and then increase in April.

**Fig. 5.**
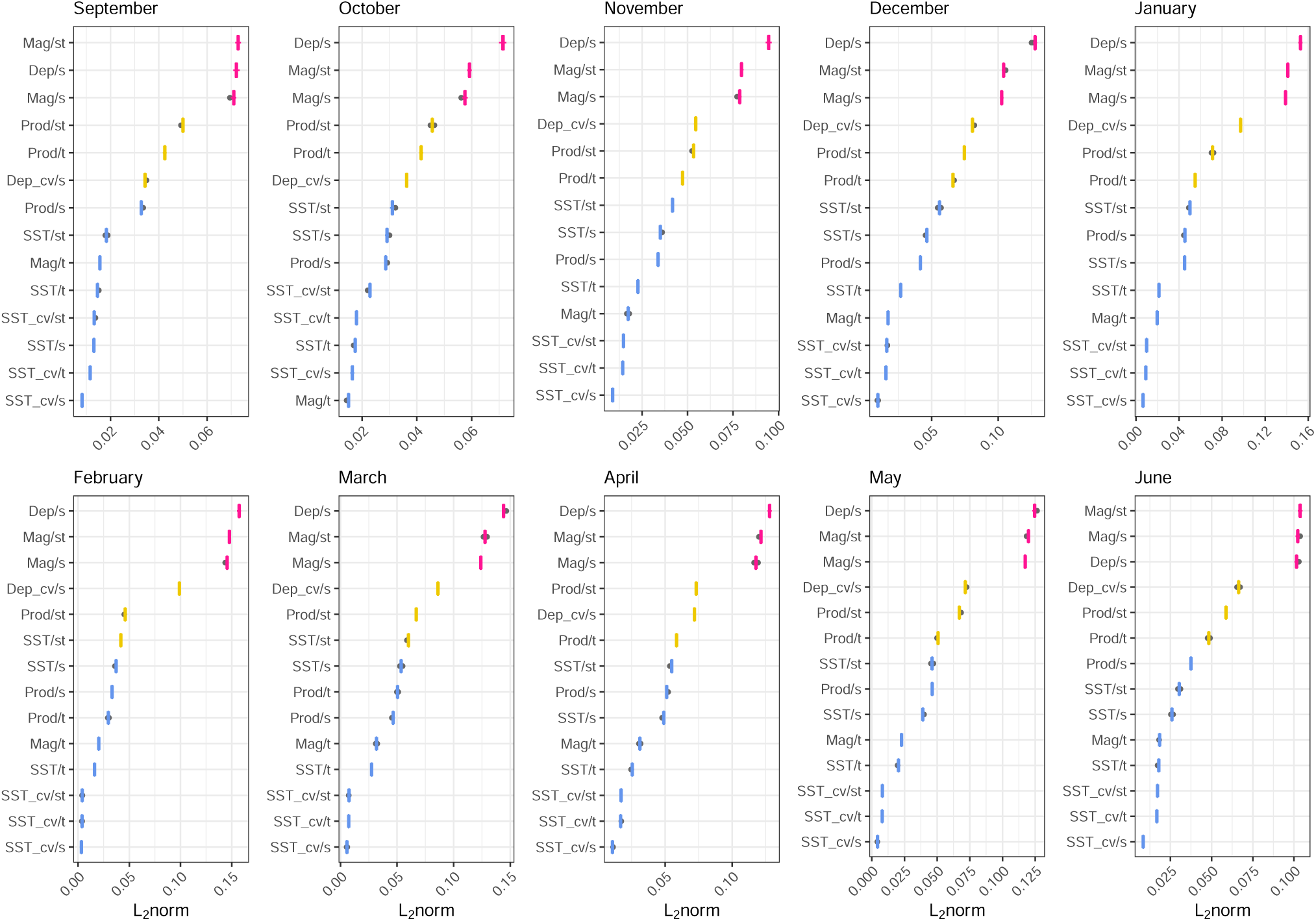
Spatial, temporal, and spatio-temporal effects of driving factors from September to June estimated from GPFI. Boxplots show the medians and the 25% and 75% quantiles. Whiskers indicate the data points within × 1.5 of the quartiles and gray circles represent outliers. Large *L*_2_ norm indicates more important factors for predicting the estimated occurrence rates of spiny dogfish. Letters after the slash (/) on the y-axis indicate spatial (s), temporal (t), and spatio-temporal (st) effects. *Dep, Mag, Prod*, and *SST* are depth, magnetic fields, productivity, and monthly mean SST, respectively. *Dep_cv* and *SST_cv* are the coefficient of variation of depth around the estimated points (i.e., submarine topography) and monthly coefficient of variation of SST (i.e., monthly variability of SST), respectively. A detailed description of parameters is provided in *2*.*1*.*2 Data collection for candidate driving factors*

The factors with weak effects were consistent across months (Fig. 5). The temporal effect of magnetic fields was always weak, although the spatial and spatio-temporal effects of these factors were always large. Similarly, the temporal effect of SST was always weak, although the spatial and spatio-temporal effects of SST changed seasonally. Effects of the CV of SST were consistently weak.

## 4 Discussion

The migration timing of spiny dogfish advanced by a month after 2000 (Fig. 4), whereas the geographic locations of migration have not changed over 5 decades (Figs. 2-3). Seasonal changes in driving factors involved changes mainly in the importance of SST; spatial and spatio-temporal effects of SST tended to increase in February (when spiny dogfish migrate southward) and decrease in April (when the species migrates northward), although temporal effects of SST were always weak (Fig. 5). These results provide two key insights. (i) The advanced timing of migration in spiny dogfish is likely related to SST. (ii) Migration is not directly driven by SST (i.e., it is not merely a reaction to changes in specific conditions) but reflects tracking of areas with the preferred SST (i.e., actively seeking specific conditions), as evidenced by the weak temporal effect of SST and elevated spatial and spatio-temporal effects of SST during southward migration. In the study area, the mean SST increased, particularly after 2000 (Fig. S1). Therefore, our results imply that spiny dogfish actively seek preferential SST during the peak period of migration for parturition; increases in SST associated with global warning have resulted in a shift to earlier migration in the species.

Marine species are more sensitive to climate change compared with species on land (e.g., Sunday et al. 2012, Pinsky et al. 2019), and disruptions in their natural phenology and distribution have been reported (e.g., Parmesan and Yohe 2003, Edward and Richardson 2004, Perry et al. 2005, Pinsky et al. 2013, Poloczanska et al. 2013, Asch 2015, Brown et al. 2016, Kanamori et al. 2019). Distributional shifts are thought to occur by tracking climate velocity (the rate and direction of the isotherms shifts through space) (e.g., Burrows et al. 2011, Pinsky et al. 2013, Sunday et al. 2015) whereas phenological shifts, which are relatively less reports than distributional shifts in marine systems, are thought to occur by earlier warming (e.g., Edward and Rechardson 2004, Asch 2015). Despite warming in the present study area, a distributional shift and an effect of temporal changes in SST were not detected.

Alternatively, our results indicated that spiny dogfish seek their suitable SST location where SST has increased, especially after 2000, resulting in advancing their migration timing. These findings provide new evidence that phenological and distributional shifts have occurred by the combination of the way that previous studies assumed, suggesting the necessity to simultaneously evaluate phenological and distributional shifts and to understand processes of those shifts for inferences in the future.

The spatial effects of magnetic fields, depth, and CV of depth (i.e., submarine topography) and spatio-temporal effect of magnetic fields were always large (Fig. 5). Accordingly, spiny dogfish had a strong preference for geographic features and therefore the migration area remained stable over 5 decades (Figs. 2-3). However, we need to have a concern for the effects of productivity (especially spatio-temporal effect of productivity), which showed consistently large effects following the effects of magnetic fields, depth, and submarine topography. This is because that the productivity in marine systems is influenced by various factors, such as climate change (e.g., Barange et al. 2014, Free et al. 2019) and overfishing (e.g., Link and Watson 2019), leading to that changes in productivity will alter the migration of spiny dogfish.

### Utility of the spatio-temporal model and machine learning

Machine learning methods have various advantages that the models can include nonlinearity and complex interactions among covariates as well as high predictability. In contrast, the models require large datasets which are often difficult to obtain in ecology and conservation fields. We overcame this disadvantage by predicting occurrence rates in many locations using a spatio-temporal model. Our approach not only increased the data used for machine learning models but also removed noise associated with data collection. Spatio-temporal models are a well-known method for standardization, that is, removing various kinds of noise (e.g., spatio-temporal changes in survey points and survey effort), effects of target fishing, environmental factors, and species misidentification (e.g., Thorson and Barnett 2017, Thorson et al. 2017a,b, Thorson 2019, Kanamori et al. 2019, 2021). Thus, the combination of spatio-temporal models and machine learning algorithms can contribute to research on the relationships between species and external factors.

Understanding the spatio-temporal patterns of species occurrence and its process is a major goal in ecology and conservation (Picl et al. 2017). Machine learning methods have used to inference species distribution and relating environmental factors because of their high predictability (e.g., Hastie et al. 2009, Maloney et al. 2012, McLaren et al. 2018, Ryo et al. 2018, Shiferaw et al. 2019), and the interpretable methods of machine learning have developed as well (e.g., Molnar 2020, Ryo et al. 2020). However, although interpretable methods can provide the importance of factors for species distribution, it was difficult to more deeply understand how those factors affect species distribution because the data frequently include spatial and temporal aspects and we did not know how those factors are important (i.e., do those factors affect the spatial, temporal, or spatio-temporal patterns of species distribution?). In this study, we challenged to evaluate the spatial, temporal, and spatio-temporal effects of driving factors on migration by an interpretable machine learning method, resulting in we inferred the process driving long-term change in seasonal migration of spiny dogfish; migration area tends not to change by the spatial effects of geographic features (magnetic fields and depth) but migration timing has advances by seeking their suitable SST location (i.e., spatial effect) where SST has increased. This partitioning method provided more simple interpretation than coefficients of linear model with many interactions (e.g., our linear model for comparing with gradient boosting) but more detailed information than importance of previous interpretable methods. Therefore, this partitioning method can help to infer the process underlying the spatio-temporal variations of species.

### Future directions

Although phenological shifts associated with global warming have been reported in marine organisms (e.g., Edward and Rechardson 2004, Asch 2015, Kanamori et al. 2019), little is known about phenological shifts of apex predators (c.f., bluefin tunas Dufour et al. 2010, tiger shark Hammerschlag et al. 2022). Previous studies have revealed the disruption of communities and ecosystems after decreases in the abundance of sharks (e.g., Dulvy et al. 2000, Heithaus et al. 2008, Polovina et al. 2009, Heithaus et al. 2012, Hummerschlag et al. 2019). In addition, the migration of apex predators contributes to marine ecosystems by horizontally and temporally connecting habitats and trophic levels (e.g., Dulvy et al. 2000, Heithaus et al. 2008, Polovina et al. 2009, Heithaus et al. 2012, Hummerschlag et al. 2019). Thus, studies of the effects of phenological shifts of apex predators on communities and ecosystems under climate change are required.

Information about population ranges and movement patterns between populations is necessary for management (Frisk et al. 2014, Crossin et al. 2017). We found areas of high species occurrence not only in NP but also in SJ, and these were connected through the Tsugaru Straits (Figs. 2 and S2-7). These results suggest that spiny dogfish can be considered a single population. Stock assessments of spiny dogfish in Japan have been conducted separately in a part of NP and in the Tsugaru Straits (Fig. 1); therefore, individuals in SJ have not been included. Thus, it is necessary to determine how the spatial range influences current stock assessment results, including comparisons of results obtained when treating the species as a single population around Japan, which should improve the stock assessment and management of spiny dogfish.

The occurrence rate of spiny dogfish increased everywhere within the study area from January to February, and then decreased gradually in the whole the study area even though the occurrences rate of spiny dogfish were moderately high levels on the north side of the Shimokita Peninsula (Figs. 2 and S2-7). These results suggest that some individuals use this area seasonally and others stay in the area year-round. In sharks, movements and habitats generally differ between sexes and life stages; females migrate from adult habitats, including migration from the mating site to parturition site, and juveniles stay in the juvenile habitat (Chapman et al. 2015). In spiny dogfish, they are thought to migrate southward for parturition (Kojima 1958) and to migrate northward for feeding (Yusa and Ishida 1956), although Ukawa (1955) hypothesized that small individuals stay throughout the year. Yano et al. (2017) recently showed that male individuals tended to be caught more frequently than female individuals on the north side of Shimokita Peninsula from 2007 to 2011, and male individuals were identified as mature based on the length-maturity relationship (Yamamoto and Kibesaki 1950). Therefore, individuals who migrate seasonally and use this area year-round may be mainly females and residents may be males. Given that sharks have long generation times and a low intrinsic population rate due to slow growth, late maturation, and low young production, population recovery after depletion is difficult (Dulvy et al. 2017, Pacoureau et al. 2021). In addition, sharks are thought to return their natal site, because there are observations that they often collapse by fishing at local geographic scales and it is difficult to recover population quickly by immigration (Chapman et al. 2015). Indeed, we found that strong preference for geographic features. Thus, both collecting individual-level data (e.g., sex, body size, and maturity) and developing management strategies, such as not only how much we can catch in a year but also when and where we can fish in a year, are necessary for sustainable fishing of spiny dogfish.

## Conclusion

This study revealed that the geographic location of migration of spiny dogfish has not changed over ca. 5 decades, whereas the timing of migration has advanced by a month after 2000 in the western North Pacific. The spatial and spatio-temporal effects of magnetic fields and depth were consistently large and the spatial and spatio–temporal effects of SST increased in the timing of migration, even though temporal effect of SST was consistently low. These results reveal that the migration area of spiny dogfish has been stable because it is determined by geographic features, whereas the migration timing has advanced. This advancing of migration timing was not a direct response to temporal changes in SST but a consequence of tracking a suitable SST location where SST increased steeply after 2000. Therefore, not only temperature but also other many factors influence migration simultaneously under climate change, underlining the importance of paying attention biotic/abiotic factors including temperature and process-based understanding to predict future impacts of climate change on phenological shifts.

## Acknowledgments

We appreciate Ms. Akane Yoshikawa, Fisheries Resources Institute, Japan Fisheries Research and Education Agency for organizing the fisheries statistics in the Sea of Japan. This research was financially supported by Japanese Ministry of Agriculture, Forestry and Fisheries, and Japan Sea National Fisheries Institute.

## Authorship

YK conceived of the research idea. TY and YY organized the catch statistics. YK and HO designed statistical analyses, and YK wrote programs and performed the analyses. YK also wrote the manuscript with input from all co-authors’ comments.

## Literature cited

Ackerman JT, Kondratieff MC, Matern SA, Cech JJJ (2000) Tidal influence on spatial dynamics of leopard sharks, Triakis semifasciata, in Tomales Bay, California. Environ Biol Fishes 58:33–43

Anderson JM, Clegg TM, Véras LVMVQ, Holland KN (2017) Insight into shark magnetic field perception from empirical observations. Sci Rep 7:11042

Araújo MB, Anderson RP, Barbosa AM, Beale CM, Dormann CF et al. (2019) Standards for distribution models in biodiversity assessments. Sci Adv 5: eaat4858

Asch RG (2015) Climate change and decadal shifts in the phenology of larval fishes in the California Current eco-system. Proc Natl Acad Sci USA 112:E4065–E4074

Asch RG (2015) Climate change and decadal shifts in the phenology of larval fishes in the California Current eco-system. Proc Natl Acad Sci USA 112:E4065 – E4074

Bakka H, Vanhatalo J, Illian JB, Simpson D, Rue H (2019) Non-stationary Gaussian models with physical barriers. Spatial Statistics 29:268–288

Bakka H, Rue H, Fuglstad G-A, Riebler A, Bolin D et al. (2018) Spatial modelling with R-INLA: A review. Computational Statistics 10:e1443

Barange M, Coetzee JC, Twatwa NM (2005) Strategies of space occupation by anchovy and sardine in the southern Benfuela: the role of stock size and intra-species competition. ICES J Mar Sci 62:645–654

Bauer S, Hoye J (2014) Migratory animals couple biodiversity and ecosystem functioning worldwide. Science 344:1242552

Bett NN, Hinch SG, Kaukinen KH, Li S, Miller KM (2018) Olfactory gene expression in migrating adult sockeye salmon Oncorhynchus nerka. J Fish Biol 92:2029–2038

Bett NN, Hinch SG (2015) Olfactory navigation during spawning migrations: a review and introduction of the Hierarchical Navigation Hypothesis. Biol Rev 91:728–759

Bi R, Jiao Y, Bakka H, Browder JA (2020) Long-term climate ocean oscillations inform seabird by catch from pelagic logline fishery. ICES J Mar Sci 77:668–679

Cain SD, Boles LC, Wang JH, Lohmann KJ (2005) Magnetic orientation and navigation in marine turtles, lobsters, and molluscs: concepts and conundrums. Integr Comp Biol 45:539–546

Carlisle AB, Starr RM (2009) Habitat use, residency, and seasonal distribution of female leopard sharks Triakis semifasciata in Elkhorn Slough, California. Mar Ecol Prog Ser 380:213–228

Chin A, Kyne PM, Walker TI, McAuley RB (2010) An integrated risk assessment for climate change: analysing the vulnerability of sharks and rays on Australia’sGreat Barrier Reef. Glob Change Biol 16:1936–1953

Crossin GT, Cooke SJ, Goldbogen JA, Phillips RA (2014) Tracking fitness in marine vertebrates: current knowledge and opportunities for future research. Mar Ecol Prog Ser 496:1–17

Diebel CE, Proksch R, Green CR, Neilson P, Walker MM (2000) Magnetite defines a vertebrate magnetoreceptor. Nature 406: 299–302

Dulvy et al. 2021 current biol

Dulvy NK, Fowler SL, Musick JA, Cavanagh RD, Kyne PM et al. (2014) Extinction risk and conservation of the world ‘s sharks and rays. eLife 3:e00590

Dulvy NK, Metcalfe JD, Glanville J, Pawson MG, Reynolds JD (2000) Fishery stability, local extinctions, and shifts in community structure in skates. Conserv Biol 14:283–293

Durif CM, Stockhausen HH, Skiftesvik AB, Cresci A, Nyqvist D, Browman HI (2021) A unifying hypothesis for the spawning migrations of temperate anguillid eels. Fish Fish 0:1–18

Ebert DA (2003) Sharks, rays and chimaeras of California. California Natural History Guides No. 71. University of California Press. 284p

Edwards M, Richardson A (2004) Impacts of climate change on marine pelagic phenology and trophic mismatch. Nature 430:881–884

Free CM, Thorson JT, Pinsky ML, Oken KL, Wiedenmann J, Jensen OP (2019) Impacts of historical warming on marine fisheries production. Science 363:979–983

Frisk MG, Jordaan A, Miller TJ (2014) Moving beyond the current paradigm in marine population connectivity: Are adults the missing link? Fish Fish 15:242–54

Gallagher AJ, Hammerschlag N (2011) Global shark currency: the distribution, frequency, and economic value of shark ecotourism. Curr Issues Tour 14:797–812

Guisan A, Lehmann A, Ferrier S, Austin M, Overton JMC et al. (2006) Making better biogeographical predictions of species ‘distributions. J Appl Ecol 43:386–392

Hammerschlag N, McDonnell LH, Rider MJ, Street GM, Hazen EL et al. (2022) Ocean warming alters the distributional range, migratory timing, and spatial protections of an apex predator, the tiger shark (Galeocerdo cuvier). Glob Change Biol 00:1–16

Hammerschlag N, Schmitz OJ, Flecker AS, Lafferty KD, Sih A et al. (2019) Ecosystem function and services of aquatic predators in the Anthropocene. Trends Ecol Evol 34:369–383

Hastie T, Tibshirani R, Friedman J (2009) The Elements of Statistical Learning: Data Mining, Inference, and Prediction, 2nd edn. Springer, New York

Heithaus MR, Wirsing AJ, Dill LM (2012) The ecological importance of intact top-predator populations: a synthesis of 15 years of research in a seagrass ecosystem. Mar Freshw Res 63:1039–1050

Heithaus MR, Frid A, Wirsing AJ, Worm B (2008) Predicting ecological consequences of marine top predator declines. Trends Ecol Evol 23:202–210

Heupel M, Simpfendorfer C, Espinoza M, Smoothey A, Tobin A, Peddemors V (2015) Conservation challenges of sharks with continental scale migrations. Front Mar Sci 2:1–7

Hoegh-Guldberg O, Bruno JF (2010) The impact of climate change on the world’s marine ecosystems. Science 328:1523–1528

Kalmijn AJ (1982) Electric and magnetic field detection in elasmobranch fishes. Science 218:916–918

Kanamori Y, Nishijima S, Okamura H, Yukimi R, Watai M, Takasuka A (2021) Spatio-temporal model reduces species misidentification bias of spawning eggs in stock assessment of spotted mackerel in the western North Pacific. Fish Res 236:105825

Kanamori, Y., Takasuka, A., Nishijima, S., Okamura, H. 2019. Climate change shifts the spawning ground northward and extends the spawning period of chub mackerel in the western North Pacific. Mar. Ecol. Prog. Ser. 624:155–166.

Keller BA, Putman NF, Grubbs RD, Portnoy DS, Murphy TP (2021) Map-like use of Earth’s magnetic field in sharks. Current Biol 31:1–6

Koch AL, Carr A, Ehrenfeld DW (1969) The problem of open sea navigation: the migration of the green turtle to Ascension Island. J Theor Biol 22:163–179

Kojima I (1958) North Pacific spiny dog fish. Tsushima Danryu Kaihatsu Chosa Hokokusho 4:34–44 (In Japanese)

Kuhn M (2020) caret: Classification and Regression Training. https://CRAN.R-project.org/package=caret

Kuhn M (2008) Building predictive models in R using the caret package. J Statistical Software 28:1–26

La Sorte FA, Graham CH (2021) Phenological synchronization of seasonal bird migration with vegetation greenness across dietary duilds. J Anim Ecol 90:343–355

Link JS, Watson RA (2019) Global ecosystem overfishing: Clear delineation within real limits to production.

Marra PP, Francis CM, Mulvihill RS, Moore FR (2005) The influence of climate on the timing and rate of spring bird migration. Oecologia 142:307–315

Mathot KJ, Smith BD, Elner RW (2007) Latitudinal clines in food distribution correlate with difierential migration in the western sandpiper. Ecology 88:781–791

McLarenJD, Buler JJ, Schreckengost T, Smolinsky JA, Boone M et al. (2018) Artificial light at night confounds broad-scale habitat use by migrating birds. Ecol Lett 21:356–364

Mecklenburg CW, Lynghammar A, Johannesen E, Byrkjedal I, Christiansen JS et al. (2018) Marine fishes of the Arctic Region.Volume I. Conservation of Arctic Flora and Fauna, Akureyri, Iceland 28:1–454

Meyer CG, Holland KN, Papastamatiou YP (2005) Sharks can detect changes in the geomagnetic field. J R Soc Interface 2:129–130

Milner-Gulland EJ, Fryxell JM, Sinclair ARE (2011) Animal Migration: A Synthesis. Oxford University Press, Oxford

Molnar C (2020) Interpretable machine learning: A guide for making black box models explainable. Lulu. com. https://christophm.github.io/interpretable-ml-book/

Nakayama SI, Fukuda H, Nakatsuka S (2020) Model-free time series analysis detected the contributions of middle-age spawner biomass and the environment on Pacific bluefin tuna recruitment. ICES J Mar Sci 77:1480–1491

Newton KC, Kajiura SM (2020) The yellow stingray (Urobatis jamaicensis) can use magnetic field polarity to orient in space and solve a maze. Mar Biol 167:36

Niella Y, Butcher P, Holmes B, Barnett A, Harcourt R (2021) Forecasting intraspecific changes in distribution of a wide-ranging marine predator under climate change. Oecologia 1–14

Orlov AM, Savinykh VF, Kulish EF, Pelenev DV (2012) New data on the distribution and size composition of the North Pacific spiny spiny dogfish Squalus Suckleyi (Girard, 1854). Scientia Marina 76:111–122

Ortega LA, Heupel MR, Van Beynen P, Motta PJ (2009) Movement patterns and water quality preferences of juvenile bull sharks (Carcharhinus leucas) in a Florida estuary. Environ Biol Fishes 84:361–373

Pacoureau N, Rigby CL, Kyne PM, Sherley RB, Winker H et al. (2021) Half a century of global decline in oceanic sharks and rays. Nature 589:567–571

Pecl GT, Araújo BA, Bell JD, Blanchard J, Bonebrake TC et al. (2017) Biodiversity redistribution under climate change: Impacts on ecosystems and human well-being. Science 355:1389

Pinsky ML, Selden RL, Kitchel ZJ (2020) Climate-driven shifts in marine species ranges: Scaling from organisms to communities. Ann Rev Mar Science 12:153–179

Pinsky ML, Worm B, Fogarty MJ, Sarmiento JL, Levin SA (2013) Marine taxa track local climate velocities. Science 341:1239–1242

Pinsky ML, Fogarty M (2012) Lagged social-ecological responses to climate and range shifts in fisheries. Climate Change 115:883–891

Polovina JJ, Abecassis M, Howell EA, Woodworth P (2009) Increases in the relative abundance of mid-trophic level fishes concurrent with declines in apex predators in the subtropical North Pacific, 1996-2006. Fish Bull 107:523–531

R Development Core Team (2020) R: a language and environment for statistical computing. R Foundation for Statistical Computing, Vienna

Rue H, Martino S, Chopin N (2009) Approximate Bayesian inference for latent Gaussian models by using integrated nested Laplace approximations. J R Statist Soc B 71:319–392

Ryo M, Angelov B, Mommola S, Kass JM, Benito BM, Hartig F (2020) Explainable artificial intelligence enhances the ecological interpretability of black-box species distribution models. Ecography 44:199–205

Ryo M, Harvey E, Robinson CT, Altermatt F (2018) Nonlinear higher order abiotic interactions explain riverine biodiversity. J Biogeogr 45:628–639

Sala OE, Chapin FS III, Armesto JJ, Berlow E, Bloomfield J et al. (2000) Global biodiversity scenarios for the year 2100. Science 287:1770–1774

Satterthwaite WH, Andrews KS, Burke BJ, Gosselin JL, Greene CM et al. (2020) Ecological thresholds in forecast performance for key United States West Coast Chinook salmon stocks. ICES J Mar Sci 77:1503–1515

Schlaff AM, Heupel MR, Simpfendorfer CA (2014) Influence of environmental factors on shark and ray mobement, behaviour and habitat use: a review. Rev Fish Biol Fisheries 24:1089–1103

Sherrill-Mix SA, James MC, Myers RA (2008) Migration cues and timing in leatherback sea turtles. Behav Ecol 19:231–236

Shiferaw H, Bewket W, Eckert S (2019) Performances of machine learning algorithms for mapping fractional cover of an invasive plant species in a dryland ecosystem. Ecol Evol 9:2562–2574

Sugihara G, May R, Ye H, Hsieh C, Deyle E, Fogarty M, Munch S (2012) Detecting causality in complex ecosystems. Science 338:496–500

Sunday JM, Pecl GT, Frusher S, Hobday AJ, Hill N, et al. (2015) Species traits and climate velocity explain geographic range shifts in an ocean-warming hotspot. Ecol Lett 18:944–953

Teitelbaum CS, Mueller T (2019) Beyond migraion: Causes and consequences of nomadic animal movements. TREE 34:569–581

Thackeray SJ, Jones ID, Maberly SC (2008) Long-term change in the phenology of spring phytoplankton: species-specific responses to nutrient enrichment and climate change. J Ecol 96:523 – 535

Thorson JT (2019) Guidance for decisions using the Vector Autoregressive Spatio-Temporal (VAST) package in stock, ecosystem, habitat and climate assessments. Fish Res 210:143–161

Thorson JT, Barnett LAK (2017) Comparing estimates of abundance trends and distribution shifts using single- and multispecies models of fishes and biogenic habitat. ICES J Mar Sci 74:1311–1321

Thorson JT, Fonner R, Haltuch MA, Ono K, Winker H (2017a) Accounting for spatiotemporal variation and fisher targeting when estimating abundance from multispecies fishery data. Can J Fish Aquat Sci 74:1794–1807

Thorson JT, Ianelli JN, Kotwicki S (2017b) The relative influence of temperature and size-structure on fish distribution shifts: a case-study on walleye pollock in the Bering Sea. Fish Fish 18:1073–1084

Thorson JT, Fonner R, Haltuch MA, Ono K, Winker H (2016) Accounting for spatio-temporal variation and fisher targeting when estimating abundance from multispecies fishery data. Canadian J Fish Aquat Scien 74:1794–1807

Torres LG, Heithaus MR, Delius B (2006) Influence of teleost abundance on the distribution and abundance of sharks in Florida Bay, USA. Hydrobiologia 569:449–455

Ubeda AJ, Simpfendorfer CA, Heupel MR (2009) Movements of bonnetheads, Sphyrna tiburo, as a response to salinity change in a Florida estuary. Environ Biol Fishes 84:293–303

Walther GR (2010) Community and ecosystem responses to recent climate change. Proc R Soc B 365:2019–2024

Walther GR, Post E, Convey P, Menzel A and others (2002) Ecological responses to recent climate change. Nature 416:389–395

Wiltschko W, Wiltschko R (2005) Magnetic orientation and magnetoreception in birds and other animals. J Comp Physiol A 191:675–693

Wiltschko R, Wiltschko W (1995) Magnetic orientation in animals. Berlin, Germany: Springer

Worm B, Davis B, Kettemer L, Ward-Paige CA, Chapman DD, et al. (2013) Global catches, exploitation rates, and rebuilding options for sharks. Mar Policy 40:194–204

Yamamoto T, Kibesaki O (1950) Studies on the spiny dog fish Squalus suckleyi. (I) on the development and maturity of the genital glands with growth. Nippon Suisan Gakkaishi 15:531–538 (in Japanese with English abstract)

Yano T, Hattori T, Shibata Y, Tanaka S (2022) Over 120 years of landing trends in Japan, for the commercially exploited shark species, Squalus suckleyi. Fish Res 249:106257

Yano T, Ohshimo S, Kanaiwa M, Hattori T, Fukuwaka M et al. (2017a) Spatial distribution analysis of the North Paciffc spiny spiny dogfish, Squalus suckleyi, in the North Pacific using generalized additive models. Fish Oceanogr 26:668–679

Yano T, Hattori T, Tamukai T, Ohshimo S (2017b) Body-length frequency and spatial segregation of the North Pacific spiny dogfish Squalus suckeleyi in Tsugaru Strait, northern Japan. Fish Sci 83:917–928

Ye H, Beamish RJ, Glaser SM, Grant SCH, Hsieh CH et al. (2015) Equation-free mechanistic ecosystem forecasting using empirical dynamic modeling. E1569–E1576

